# Mime: A flexible machine-learning framework to construct and visualize models for clinical characteristics prediction and feature selection

**DOI:** 10.1101/2023.11.28.569007

**Authors:** Hongwei Liu, Wei Zhang, Yihao Zhang, Abraham Ayodeji Adegboro, Luohuan Dai, Zhouyang Pan, Wang Li, Kang Peng, Deborah Oluwatosin Fasoranti, Siyi Wanggou, Xuejun Li

**Author notes:** Correspondence: Siyi Wanggou, Department of Neurosurgery, Xiangya Hospital, Central South University, Changsha, Hunan 410008, China., Xuejun Li, Department of Neurosurgery, Xiangya Hospital, Central South University, Changsha, Hunan 410008, China. Hongwei Liu, Wei Zhang and Yihao Zhang have contributed equally to this work and share first authorship.

## Abstract

With the widespread use of high-throughput sequencing technologies, understanding biology and cancer heterogeneity has been revolutionized. Recently, several machine-learning models based on transcriptional data have been developed to accurately predict patient’s outcome and clinical response. However, an open-source R package covering state-of-the-art machine learning algorithms for user-friendly access has yet to be developed. Thus, we proposed a flexible computational framework to construct machine learning-based integration model with elegant performance (Mime). Mime streamlined the process of developing predictive models with high accuracy, leveraging complex datasets to identify critical genes associated with prognosis. An in silico combined model based on de novo PIEZO1-associated signatures constructed by Mime demonstrated high accuracy in predicting outcomes of patients compared with other published models. In addition, PIEZO1-associated signatures could also precisely infer immunotherapy response by applying different algorithms in Mime. Finally, SDC1 selected from PIEZO1-associated signatures presented high-potential role in glioma with targeted prospect. Taken together, our package provides a user-friendly solution for constructing machine learning-based integration models and will be greatly expanded to provide valuable insights into current fields.

## INTRODUCTION

The widespread use of high-throughput sequencing technologies has a profound impact on the understanding of biology and cancer heterogeneity[1]. An increasing number of researchers have identified specific molecular features associated with disease progression, patient outcomes and therapeutic response from sequencing data[2, 3]. These selective signatures provide a comprehensive overview of particular biological process that regulate transcriptional networks of cancer cells[4]. Due to the diversity and vastness of features, rational computational strategies are urgently needed to identify critical genes for disease.

Machine learning (ML), a branch of computer science that learns from complex datasets to develop a predictive model with high accuracy, have become a popular tool in medical researches[5]. More recently, several diagnostic and prognostic models based on machine learning have been developed from transcriptional data for various cancer types[6–8]. Since the performance of machine learning models from large amounts of transcriptional data can vary, it is often recommended to compare the results derived from several methods and select the optimal one for further application[6, 9, 10]. In addition, given various formats of input and output and system parameters supported by different in silico approaches, such as CoxBoost[11] and superpc[12], comparative analysis can become extremely complex. Nevertheless, there is still no software covering state-of-the-art machine learning algorithms for user-friendly access. Thus, integrating transcriptional data with different machine learning algorithms will be greatly expanded to provide valuable insights into current fields.

Here, we developed Mime, an open-source R package with elegant performance which simplified procedure of constructing machine learning-based integration models from transcriptomic data. To explore predicting models and candidate genes from large-scale features in one-stop, Mime mainly provided four applications: () establishment of prognosis models by integrating 10 machine learning algorithms, () construction of binary response models by applying 7 machine learning algorithms, () core feature selection related with prognosis by 8 machine learning methods and () visualization of the performance of each model. Specifically, we used de novo PIEZO1 associated signatures identified from primary glioblastoma cells and several publicly available cohorts as an example to demonstrate the workflow of Mime in detail and show its overall capabilities.

## MATERIALS AND METHODS

### Acquisition and pre-processing of glioma cohorts

Nine glioma datasets, comprising mRNA expression profiles, corresponding clinical and genomics feature data, were acquired from publicly accessible databases. Notably, transcriptional data from the TCGA-Glioma dataset (n = 702), which encompasses the TCGA low grade glioma (LGG) and glioblastoma cohorts, were retrieved from XENA (https://xena.ucsc.edu/) in June 2023. TCGA genomic features profiles were acquired from UCSCXenaShiny[13]. Four external datasets, namely CGGA.325 (n = 325), CGGA.693 (n = 693), CGGA.1018 (n = 1018), and CGGA.array (n = 301) were obtained from the Chinese Glioma Genome Atlas (CGGA, http://www.cgga.org.cn/). Clinical and transcriptomic annotations of 168 patients who had RNA-Sequencing data for at least two time points were procured from the Glioma Longitudinal Analysis Consortium (GLASS, https://www.synapse.org/glass). Based on diagnostic time points, we divided the cohorts into two datasets: the primary glioma cohort and the recurrent glioma cohort. We also obtained GSE108474 (n = 314) and GSE16011 (n = 276) from the Gene Expression Omnibus (GEO, http://www.ncbi.nlm.nih.gov/geo). We excluded samples without complete survival information from these nine glioma cohorts. The data from the Illumina HiSeq platform was converted into the transcripts per kilobase million (TPM) format. Clean microarray data was obtained by correcting the background, performing quantile normalization, and logarithmic transformation. The TCGA-Glioma dataset was employed as the training data to screen for the best predictive model, while the remaining eight glioma cohorts served as independent validation datasets. Supplementary Table S1 offers more information.

### Acquisition and pre-processing of cohort with immune checkpoint inhibitor therapy

Eighteen cohorts of pre-treatment samples with immune checkpoint inhibitors (ICI) were collected from published studies and resources, comprising a total of 1,059 patients across eight cancer types (296 responders and 763 non-responders). To obtain an integrated dataset, the R package sva (v3.40.0) was used to remove batch effects, and samples lacking response information were excluded. The unified group was arbitrarily segregated into two groups, designated as the training dataset (70%, n=730) and the validation dataset (30%, n=312). Supplementary Table S2 offers additional data on the cohorts who received ICI therapy.

### Developing the optimal prognostic model with the 117 machine learning combinations

Here, based on the previous literature, we provided a novel machine learning framework to develop the optimal model for predicting the prognosis of the patients based on the input variables and the provided cohorts[6]. First, we conducted univariate Cox regression analysis to identify prognostic features from the input variables. Genes exhibiting a p value less than 0.05 were recognized as having prognostic significance. Second, the genes were entered into the machine learning framework titled leave-one-out cross-validation (LOOCV). This framework incorporates ten classical machine learning algorithms, e.g., random forest (RSF), elastic network (Enet), stepwise Cox, CoxBoost, partial least squares regression for Cox (plsRcox), supervised principal components (SuperPC), gradient boosting machine (GBM), survival support vector machine (Survival-SVM), Ridge, and least absolute shrinkage and selection operator (LASSO). Possible variable selection filters include Lasso, stepwise Cox, CoxBoost, and RSF, each with distinct core parameters. 117 combinations were integrated into the LOOCV framework with tenfold-cross validation to screen for a hyperparameter-tuned model. Precisely ten machine learning algorithms were utilized, each with varying parameters. The model with the mean of the highest C-index in the validation cohorts was the most valuable predictive model with the best accuracy and lower risk of overfitting.

### Comparing the optimal model with the previous published predictive models

To compare the optimal predictive signature developed through the LOOCV framework with other published predictive signatures, we gathered an exhaustive selection of prognostic signatures for glioma, encompassing LGG and glioblastoma. A total of 33, 22, and 40 different predictive signatures were identified for prognosticating the outcomes of glioma, LGG, and glioblastoma patients, respectively. The characteristics that distinguish the signature and their respective coefficients have been furnished in the R package Mime and Supplementary Table S3.

### Constructing models for predicting the binary variable with seven machine learning algorithms

Seven standard machine learning algorithms, such as Naïve Bayes (NB), AdaBoost Classification Tree (AdaBoost), CancerClass, Random Forest (RF), Boost Logistic Regression (LogiBoost), K-nearest neighbors (KNN), and Support Vector Machine (SVM), were implemented to create a model for the prediction of a binary variable[14–19]. As there were no parameters that cancerclass required, the entire training dataset was utilized to train the model. To determine the optimal model, fivefold-cross validation was performed, with each resampling repeated 10 times. The model developed by the seven algorithms that the produced best area under curve (AUC) was regarded as the optimal model for downstream analysis.

### Comparing the developed response model with other predictive models for ICI therapy

To examine the effectiveness of the binary predictive model that we created based on ICI cohorts, we collected 13 previously published ICI response signatures, such as PDL1.Sig, IMS.Sig, and TRS.Sig[20–31]. The algorithms and code script were obtained from the original studies. To compare the performance of all 13 signatures and our model, we developed a visualization function in the R package Mime. Further details on the 13 ICI signatures are available in Supplementary Table S4.

### Screening out the core variables for the prognosis with eight machine learning algorithms

Here, a novel computational framework consisting of eight machine learning algorithms was constructed to screen out the core prognostic variables based on the provided transcriptomic profile and corresponding survival information. The computational framework consisted of three steps. Firstly, the prognostic factors were identified through the univariate Cox regression analysis. Secondly, we evaluated eight machine learning algorithms related to survival analysis, comprising LASSO, Enet, Boruta, CoxBoost, RSF, XGBoost, stepwise Cox, and SVM-REF, in order to identify the most crucial features. We executed LASSO 1000 times with different seeds and determined the core features selected by LASSO constituted variables that had been selected more than 50 times. The stepwise Cox regression analysis included the direction parameters ‘forward’, ‘backward’ and ‘both’, whereas Enet involved 9 alpha parameters ranging from 0.1 to 0.9. Thus, there existed 18 distinct models for screening the fundamental characteristics using diverse parameters. Thirdly, we could select the variables that have been filtered out most frequently as core features based on their selection frequency.

### Determining PIEZO1-associated signature

RNA-seq data of primary glioblastoma cell lines G508 and G532 treated with scrambled shRNA and PIEZO1 shRNA were acquired from GSE113261. Its raw sequencing data were mapped to the hg38 reference genome through HISAT2 and StringTie to obtain raw count matrix. R package DESeq2 (v1.32.0) was used for differential gene expression analysis. Differentially expressed genes (DEGs) with log2 Fold Change > 2 (or < -2) and adjusted P-value < 0.05 were defined as significant. All up-regulated and down-regulated DEGs among G508 and G532 cell lines between scrambled and PIEZO1 shRNA were intersected respectively to identify high confidence PIEZO1-associated signature.

### Enrichment analysis

SDC1-regulated genes between high- and low-expression group in TCGA, CGGA.325, CGGA.693 and GSE16011 respectively were identified by R package limma (v3.48.0). The criteria for gene filtering were set as log2 Fold Change > 0.5 (or < -0.5) and adjusted P-value < 0.05. Then GSEA algorithm in R package clusterProfiler (v4.7.1) based on average values of log2 Fold Change in four datasets were applied to conduct Gene Ontology (GO) enrichment analysis.

### Survival analysis

Patients were divided into high and low group according to the median value of specific feature for overall survival (OS) analysis. Hazard ratios (HRs) with 95% confidence intervals (CI), log-rank P values and Kaplan-Meier curves were calculated and plotted by R package survival (v3.3-1) and survminer (v0.4.9). Multivariate Cox proportional hazard model was executed by the R packages ezcox (v1.0.2).

### Statistical analysis

All the data analysis and graph generations were completed in R (v4.1.3), Adobe Photoshop software and BioRender.com. Correlations between variables were explored using Pearson or Spearman coefficients. Continuous variables fitting a normal distribution between binary groups were compared using a T-test. Categorical variables were compared using the Chi-Squared test. P-value < 0.05 was considered statistically significant, and all statistical tests were two-sided.

## RESULTS

### Overview of Mime

The schematic diagram of Mime was depicted in Figure 1, and the technical details were provided in Methods. Basically, Mime involved three steps for user-freely analysis: data input, models construction and results visualization. The first step for users was to acquire multiple cohorts containing transcriptional sequencing data with information of survival or clinical response to therapy as well as a gene set as inputs to Mime. Then, Mime applied various machine learning algorithms to train models for predicting outcome or clinical response of patients, which could also screen core features from a large number of genes. Finally, Mime provided several graphs to help users interpret the results. In this process, the overall capacity of each model was comprehensively estimated and users could select optimal model for further analysis. Especially, Mime could compare AUC or C-index of optimal model with other established models derived from previous studies if user provided. In addition, difference of immune cell infiltration and biological enrichments between samples with high and low risk score calculated by utilizing optimal model could also be determined in Mime. All functions and instructions in detail of Mime were available in the GitHub (https://github.com/l-magnificence/Mime).

**Figure 1.**
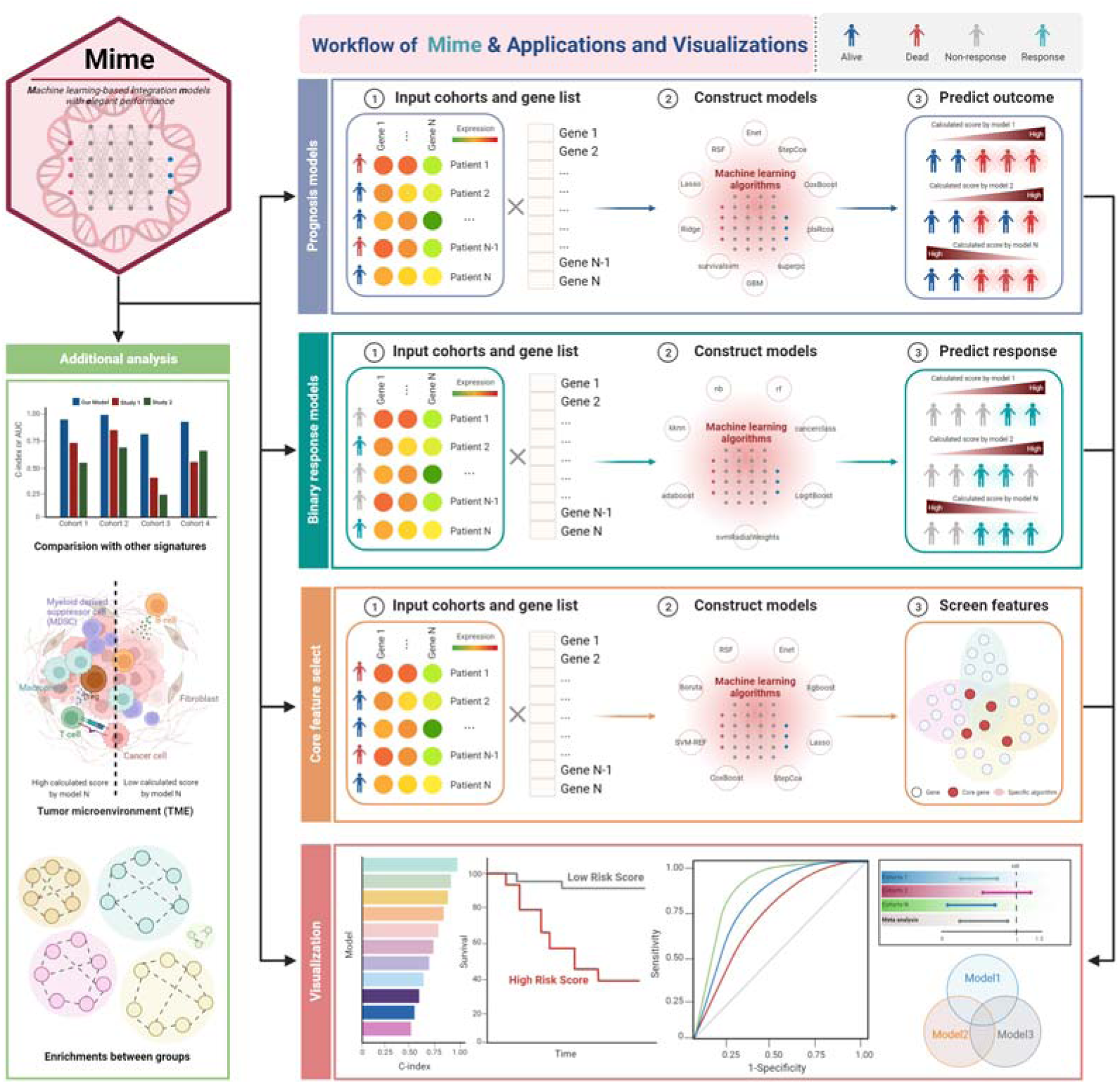
A schematic diagram of Mime. Mime streamlined the process of developing models for accurately predicting outcomes and therapeutic response of patients, leveraging complex datasets to identify critical genes associated with prognosis.

### Establishment of de novo prognosis models associated with PIEZO1 by Mime

Since PIEZO1-mediated mechanotransduction was essential for promoting glioma aggression, we used PIEZO1-asscociated signatures identified in primary glioblastoma cells from previous study as well as public glioma cohorts as an example to illustrate the application of Mime[32]. RNA sequencing was performed on primary G508 and G532 cell lines with PIEZO1 knockdown by shRNA (Figure 2A). There were totally 89 shared down-regulated genes and 38 shared up-regulated genes among G508 and G532 after PIEZO1 knockdown, which were defined as PIEZO1-associated signature (PIAS) (Figure 2B). This signature and 9 glioma transcriptomic datasets including one training cohort and eight validation cohorts were then used to construct models by integrating 10 machine learning algorithms in Mime. Among 117 models constructed by Mime, StepCox[forward]-Ridge combined model (STRICOM) had the highest mean of C-index in validation cohorts as well as in all cohorts indicating its outstanding performance (Figure 2C, Supplementary Figure S1A). Indeed, the expression level of most genes in STRICOM were decreased in PIEZO1-knockdown cells (Supplementary Figure S1B). We further separated glioma patients into high-risk and low-risk group according to the median of risk score calculated by Mime based on STRICOM and determined its survival probability in each cohort. Interestingly, patients with high risk score had significantly worse outcomes in all cohorts (Figure 2D). These results demonstrated that Mime made it easy for users to build prognostic models based on provided gene set and datasets.

**Figure 2.**
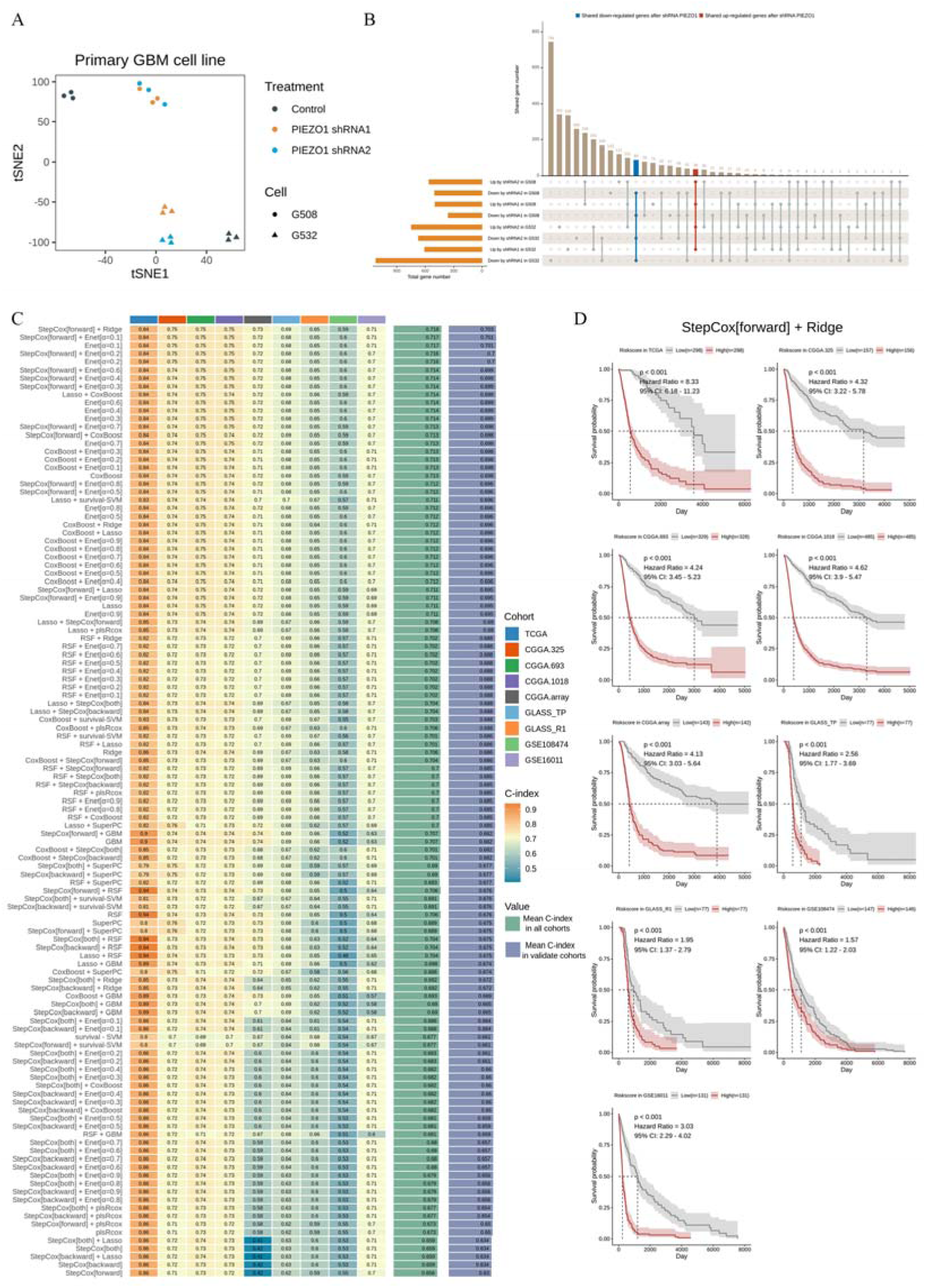
Construction of prognostic models based on PIEZO1-associated signature. **A**. The basic information about primary glioblastoma cell lines. t-SNE: t-distributed stochastic neighbor embedding. **B**. DEGs identified between control and PIEZO1 knockdown condition in G508 and G532. Left: number of DEGs in each condition. Top: number of DEGs intersected in multiple conditions. **C**. C-index of each model among different cohorts sorted by the average of C-index in validate cohorts. **D**. The relation between risk score calculated by StepCox[forward]-Ridge combined model and outcome of patients in different cohorts.

### Power evaluation of optimal model

As another metrics used to evaluate a prognostic model was AUC, we performed a time-dependent ROC curve analysis of STRICOM through Mime. Of note, the 1-year and 5-year AUC predicted by STRICOM ranked first with the highest mean of AUC in validation cohorts, although 3-year AUC were not top one among all models (Figure 3A, Supplementary Figure S1C-D). In particular, some validation cohorts such as CGGA, GLASS and GSE16011, also presented high AUC compared with TCGA training cohort predicted by STRICOM (Figure 3B, Supplementary Figure S1E). These low power of model in GSE108474 may be due to the quality of microarray data. To determine the prognostic effect of STRICOM, we performed meta-analysis of univariate COX regression via Mime, which showed that score calculated by STRICOM was the risk factor in glioma (Figure 3C). Having uncovered some known molecular biomarkers for glioma, we further performed multivariate COX regression analysis and found that score calculated by STRICOM was an independent prognostic factor taking into account gender, age at diagnosis, WHO grade, IDH mutation, 1p/19q codeletion and MGMT promoter methylation in multiple datasets (Figure 3D). Together, these results revealed that the optimal model constructed by Mime presented high accuracy in predicting outcomes of patients.

**Figure 3.**
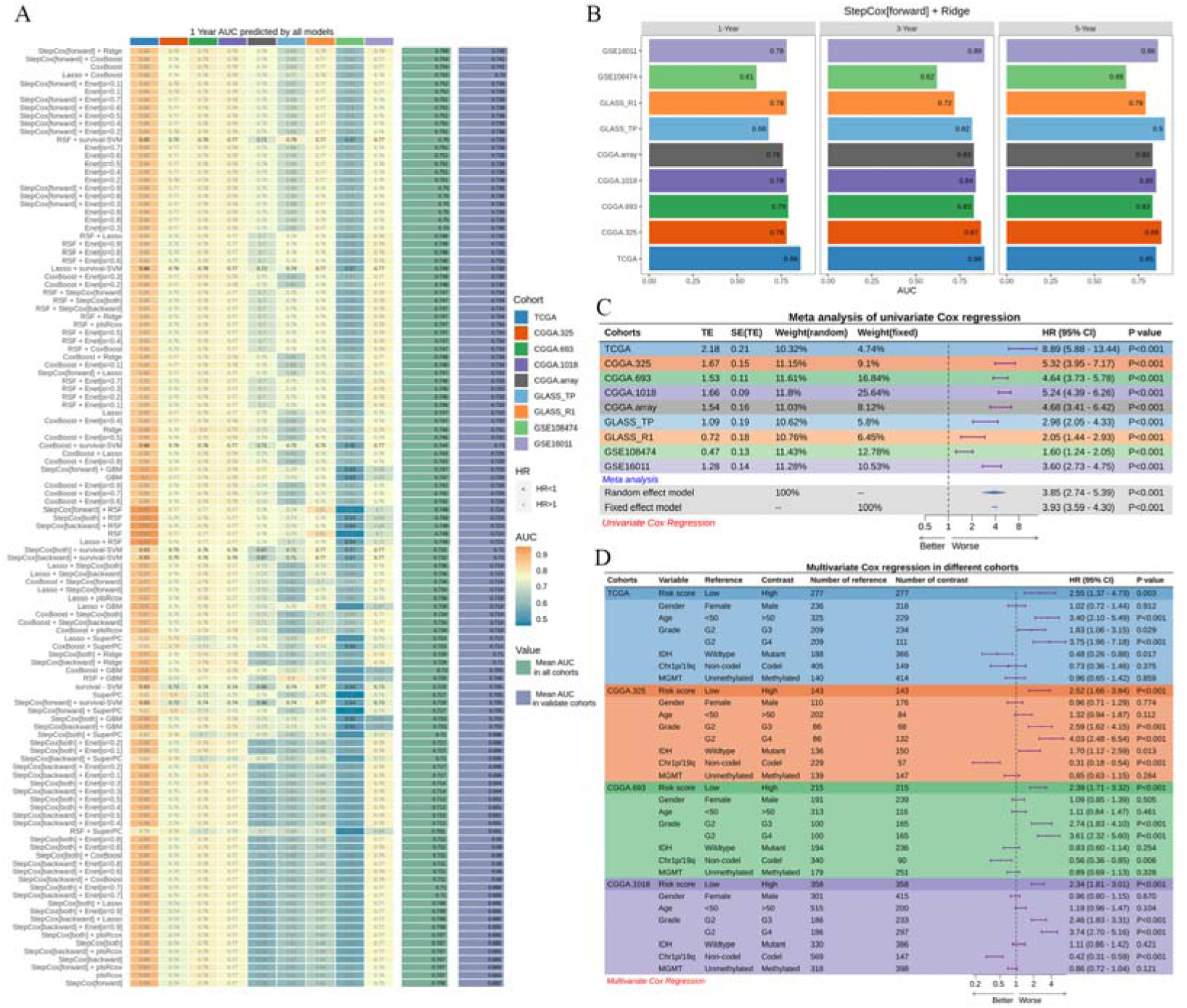
Performance of prognostic models. **A**. 1-year AUC of each model among different cohorts sorted by the average of AUC in validate cohorts. Black font of number mean that risk score calculated by this model predicted better outcome in corresponding cohort, otherwise predicted worse outcome. **B**. 1-year, 3-year and 5-year AUC of StepCox[forward]-Ridge combined model among different cohorts. **C**. Meta-analysis of univariate cox result of StepCox[forward]-Ridge combined model among different cohorts. **D**. Multivariate cox result of StepCox[forward]-Ridge combined model in four independent cohorts.

### Comparison of established models based on gene expression

Recently, a large number of prognostic and predictive models based on machine learning had been developed in glioma with the development of next-generation sequencing[33]. To comprehensively compare the performance of STRICOM with other published models in glioma, we retrieved 95 models from previous study, which were packaged in Mime. Of course, users could also provide their own models of specific disease to Mime for comparison. In our study, we performed univariate Cox regression for each model across all datasets to compare the relationship between models and prognosis and noticed that STRICOM was significantly associated with worse outcomes across all cohorts compared with other models (Figure 4A). Furthermore, STRICOM displayed more excellent performance than most models in almost all cohorts when comparing the C-index (Figure 4B). Similarly, 1-year, 3-year and 5-year AUC of STRICOM also ranked among the best across almost all cohorts compared with other models (Figure 4C, Supplementary Figure S2A-B). Collectively, these results suggested that STRICOM had better extrapolation potential and could be conveniently compared with other models in Mime.

**Figure 4.**
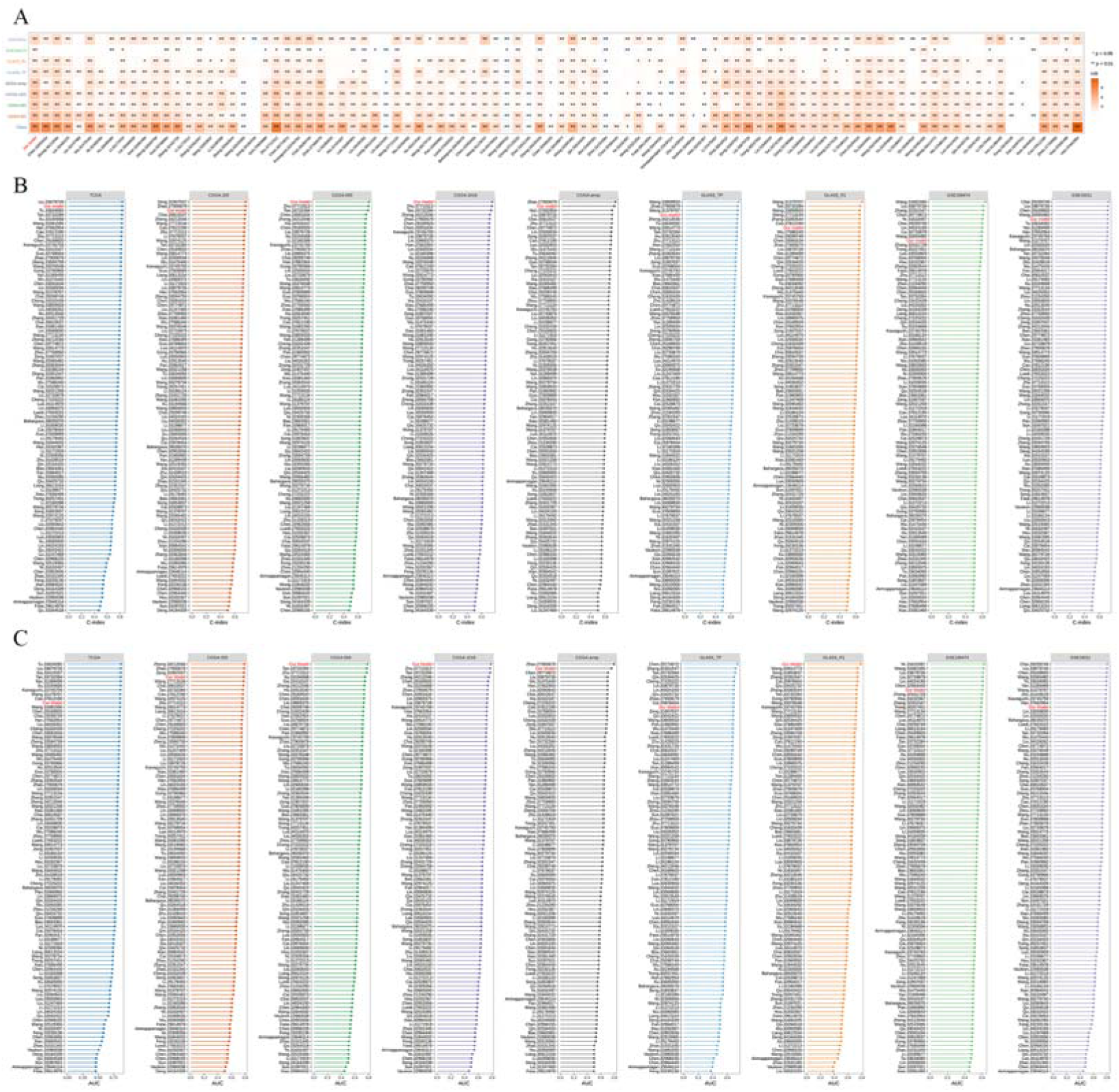
Comparison with previously established models in glioma. **A**. HR of StepCox[forward]-Ridge combined model and 95 published models across 9 cohorts. **B**. C-index of StepCox[forward]-Ridge combined model and 95 published models across 9 cohorts. **C**. 1-year AUC predicted by StepCox[forward]-Ridge combined model and 95 published models across 9 cohorts.

### Depicting the microenvironment and genome landscape shaped by STRICOM

In order to facilitate downstream analysis for users after establishing a prognostic model, Mime integrated the immune infiltration and tumor microenvironment signatures from R packages immunedeconv[34, 35] and IOBR[36], allowing users to visualize results quickly. Through tumor microenvironment analysis, we observed that, in both TCGA and CGGA cohorts, the immune infiltration scores were higher in the high-risk group compared to the low-risk group for STRICOM (Figure 5A-B). Additionally, in the TCGA-Glioma cohort, the expression of many important immune-related genes, such as CXCL10, CD276, TNFRSF14, was highly correlated with the risk score of STRICOM model (Figure 5C). Furthermore, we found a significant correlation between the high-risk group based on STRICOM and genomic features such as higher loss of heterozygosity, CNA alteration fraction, homologous recombination deficiency (HRD) score, non-silent mutations, and aneuploidy score, indicating a higher level of genomic instability in the high-risk group (Figure 5D-F). These results might explain, to some extent, why there is an apparent stratification in the prognosis of patients based on score calculated by STRICOM.

**Figure 5.**
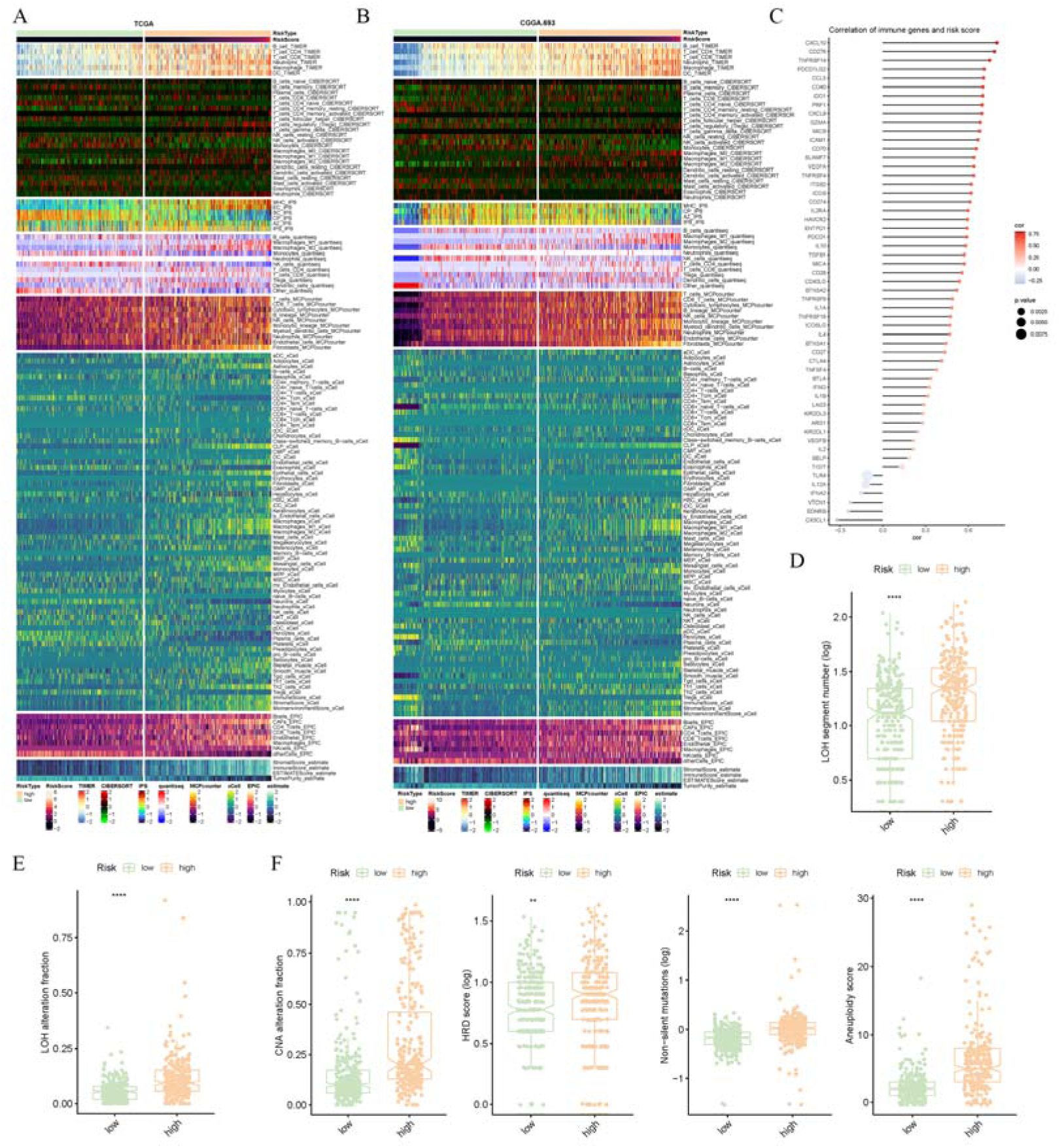
Correlation between risk score and immune or genome signatures. **A**. Relationship between risk score calculated by StepCox[forward]-Ridge combined model and microenvironment signatures deconvoluted by different methods in TCGA glioma cohort. Method IPS was from package IOBR, while others were from package immunedeconv. **B**. Same as A but in CGGA.693 cohort. **C**. Correlation between risk score and various immune genes. **D-F**. Correlation between risk score and loss of heterozygosity segment number (D), loss of heterozygosity alteration fraction (E), CNA alteration fraction, homologous recombination deficiency score, non-silent mutations, and aneuploidy score (F) respectively. \**P* < 0.05, \*\**P* < 0.01, \*\*\**P* < 0.001, \*\*\*\**P* < 0.0001.

### Development of predicting models for therapeutic response by Mime

Accumulating clinical trials of immune checkpoint inhibitor treatment had been ongoing in various cancer types, however only a relatively small proportion of patients responded to it[37]. Here, we pooled 18 cohorts in which patients received anti-PD(L)-1 or anti-CTLA4 therapy, with a total of 1042 patients divided into a training set (70%) and a validation set (30%) similar to the process of previous study[38]. Next, we provided PIAS to Mime in order to construct predicting models by using seven different machine learning algorithms (Figure 6A). Notably, the AUC of Adaptive Boosting model (adaboost) achieved 1 in training set and 0.674 in validation set, which were better than other models (Figure 6B). In addition, the performance of our developed model was more excellent than other previously published ICI related models when comparing AUC (Figure 6C). Taken together, Mime covered the main models for users to comprehensively analyze the potentials of specific signatures.

**Figure 6.**
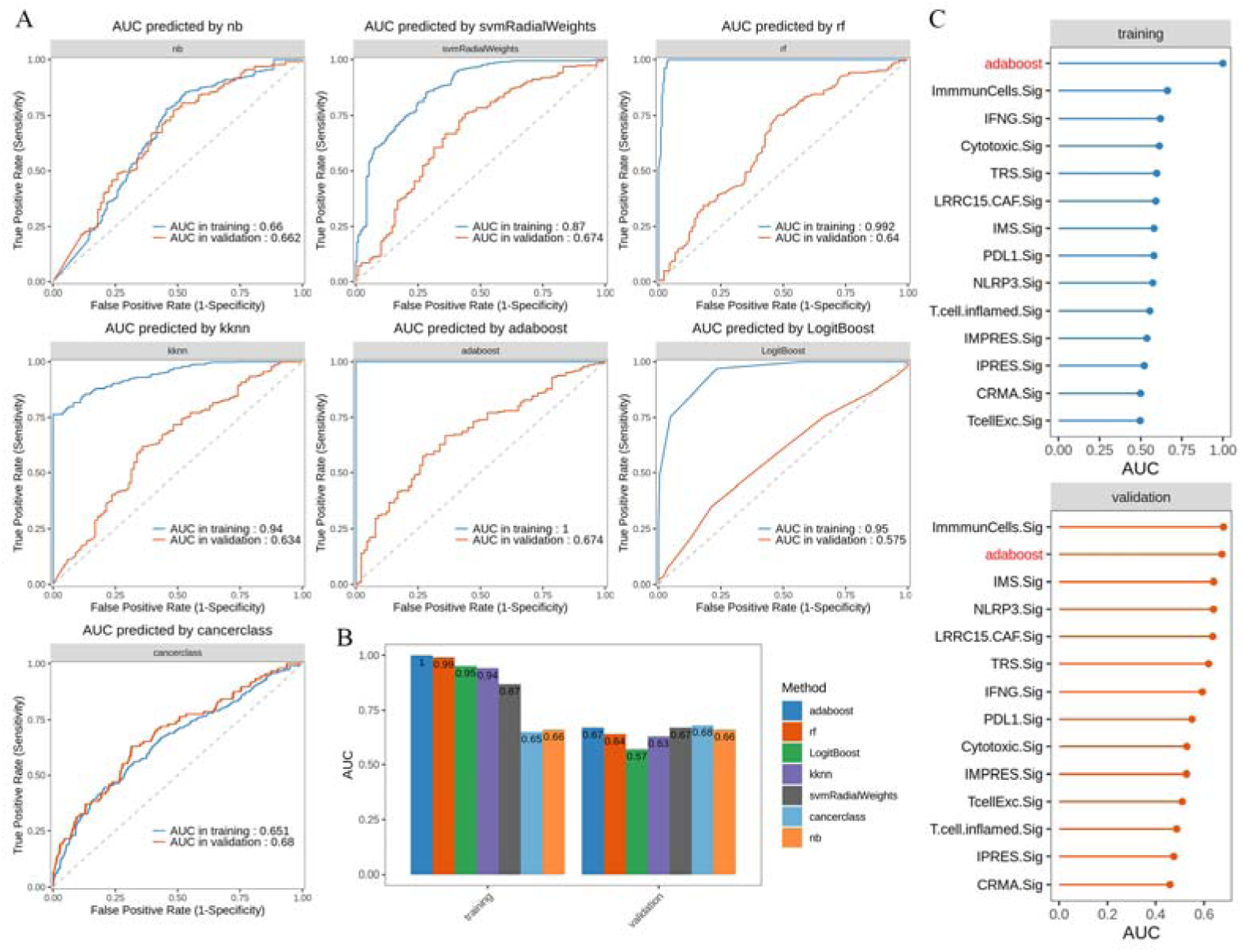
Construction of predicting models for immunotherapy benefits. **A**. ROC curves of each model to predict the benefits of immunotherapy in training and validation dataset. **B**. The distribution of AUC predicted by 7 machine-learning models in training and validation dataset. **C**. AUC predicted by adaboost model and 13 published models across training and validation dataset.

### Identification of critical gene via Mime

To investigate the potential genes for in-depth study, PIAS and TCGA glioma cohort were also provided to Mime for core feature selection by different algorithms (Figure 7A). Most top selected genes (Figure 7B), such as AQP1, TOP2A and GJB2, were well-known as crucial targets in various diseases[39–41]. Intriguingly, SDC1, a member of the syndecan proteoglycan family also named CD138, had been reported to enhance the radioresistance of glioblastoma via influencing the fusion of autophagosomes with lysosomes[42, 43]. Thus, we chose SDC1 as an example to further illustrate its potential role in glioma. Indeed, the expression of SDC1 was reduced when PIEZO1 was knocked down in both G508 and G532 (Figure 7C). Consistent with our findings, public datasets consisting of TCGA, CGGA, GLASS, GSE108474 and GSE16011, also showed a significant positive correlation between PIEZO1 and SDC1 (Figure 7D). Besides, high expression of SDC1 was significantly associated with higher grade, malignant histology, IDH wild type, 1p/19q non-co-deletion and mesenchymal subtype in glioma cohorts (Figure 7E). In order to demonstrate the involved biological mechanisms by SDC1, we separated patients into high-expression and low-expression group based on median expression level of SDC1 for identification of DEGs. Then, DEGs alone in four independent datasets presented in Figure 7E were intersected to obtain SDC1-regulated genes, which were used to perform GO enrichment analysis (Figure 7F). Totally, there were 636 shared upregulated genes and 295 shared downregulated genes in SDC1 high-expression group. Enrichment network suggested that upregulated genes were associated with regulation of cell cycle, cytoskeleton organization, extracellular matrix organization, cell-substrate adhesion and cellular response to stimulus, while downregulated genes were associated with neurotransmitter signaling and synaptic structure (Figure 7G). These changes of biological processes associated with SDC1 were consistent with the known roles of PIEZO1 in regulating glioma aggression[32]. Collectively, Mime-identified gene presented high-potential role in source disease with targeted prospect.

**Figure 7.**
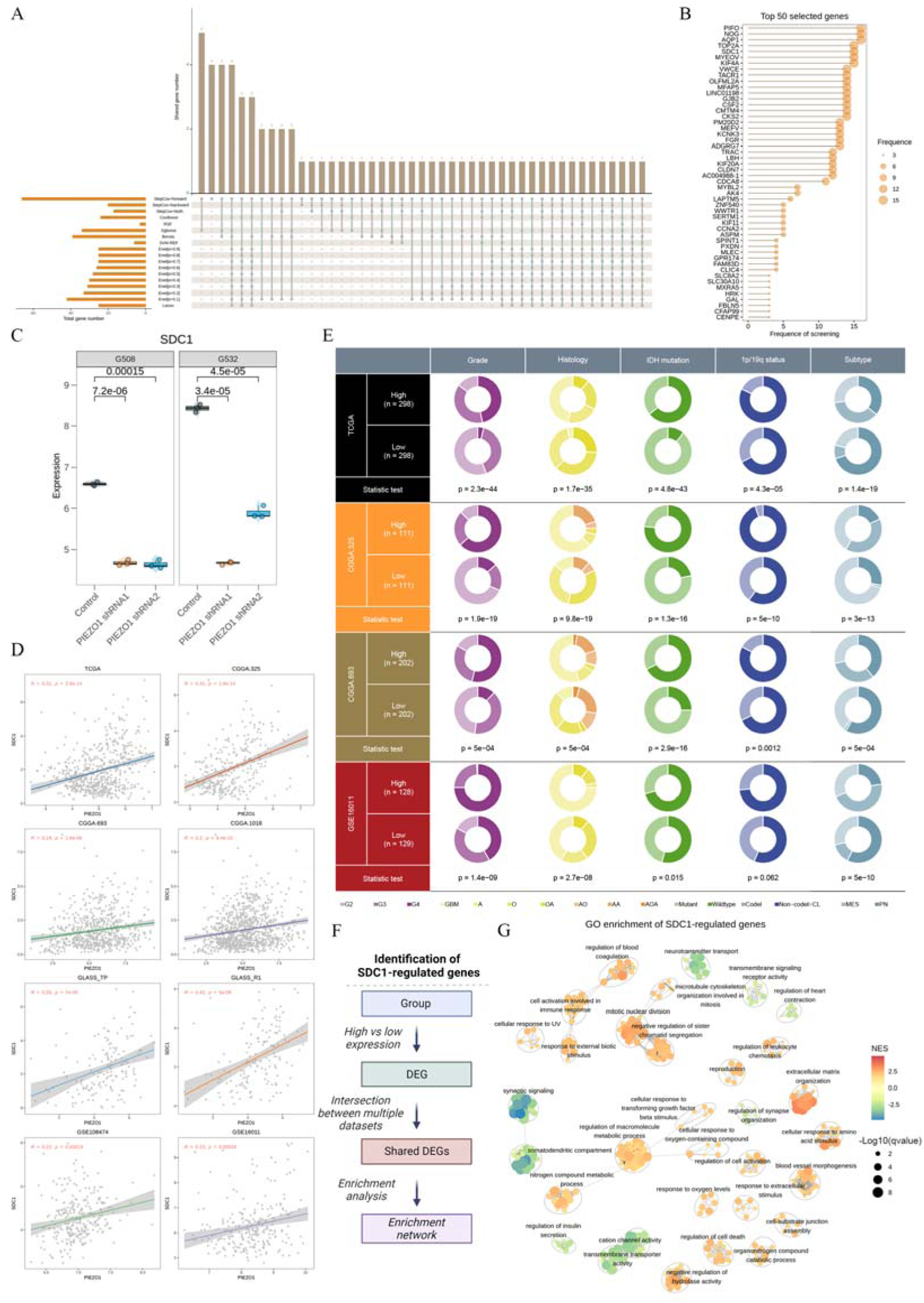
Characteristic of SDC1 in glioma. **A**. Prognosis-associated genes selected by different machine-learning algorithms. Left: number of genes filtered by each model. Top: number of genes intersected by multiple models. **B**. Frequency of genes selected by different models. **C**. Expression level of SDC1 between control and PIEZO1 knockdown condition in G508 and G532. Statistic test: t-test. **D**. Pearson correlation between PIEZO1 and SDC1 in different cohorts. **E**. The relationship between expression of SDC1 and other clinical features (Grade, Histology, IDH mutation, 1p/19q status and transcriptional subtypes) in TCGA, CGGA.325, CGGA.693 and GSE16011 cohorts. Statistic test: chi-square test. **F**. Workflow showing how to identify SDC1-regulted genes. **G**. GO enrichment of SDC1-regulted genes. Normal enrichment score (NES) > 0 indicated up-regulated processes in SDC1 high-expression group otherwise down-regulated processes in SDC1 high-expression group.

## DISCUSSION

As the application of machine learning expands across various domains such as weather prediction and recommendation engines, an increasing number of researchers recognize the potential to apply novel machine learning methods developed in other fields in medical area[44–46]. This has led to the emergence of a plethora of machine learning-based prognostic models. While researchers have the flexibility to choose a particular class of machine learning algorithm from different perspectives, the comparison and selection of the optimal model remain a complex task due to the diverse requirements in file formats, operating environments, and other factors across different machine learning methods. It is challenging for a single user to comprehensively compare the effectiveness of various algorithms on the same training dataset and validation datasets in order to obtain an advanced prognostic model.

Recently, more researchers have begun using combinations of various machine learning algorithms for more accurate and stable model construction[6–8]. To facilitate a more straightforward evaluation of the strengths and weaknesses of different models, we developed the R package Mime to simplify the process of building machine learning ensemble models from transcriptome data. As an example, we selected the PIAS in glioma for prognostic model construction. After comparing the predictive performance of 117 different machine learning models, we identified the outstanding model STRICOM. Simultaneously, we reviewed 95 glioma prognostic models published in recent years, and in comparison, STRICOM remained one of the most superior models. This example not only highlights the practicality of Mime but also underscores the tremendous potential of machine learning in prognosis research. Besides, Mime integrates core functions such as response model construction, feature selection, immune infiltration analysis and data visualization, enhancing a deep understanding of models for researchers and revealing the crucial functions these high-value potential genes may play in diseases. Based on these tentative explorations, users can further perform downstream analyses to validate corresponding biological functions and phenotypes for specific feature.

Although we use transcriptional data as an example to demonstrate the applications of Mime, it can support other input data such as proteomic data, radiomic data, clinical biological indicator data and other numerical matrices with clinical information of patients. For Mime, we also acknowledge its limitations. Firstly, Mime integrates a vast number of machine learning algorithms, and due to variations in users’ operating environments, Mime’s computational speed may be slow in certain extreme cases, with high computational resource requirements on large-scale datasets. Additionally, Mime’s performance is highly dependent on the quality and consistency of user-input data. Poor data quality or non-compliance with standards may impact the accuracy and reliability of the model. Our future research directions may include further improving Mime’s computational speed and addressing compatibility issues in complex operating scenarios. Finally, as novel machine-learning models developed in the future, we will supplement more models in the next version of Mime to meet the needs of users.

In summary, our study provides a comprehensive and powerful open-source R package, Mime, making it easier for researchers to integrate various machine learning algorithms for a better analysis of specific signatures. We hope that Mime can provide more researchers with stable, reliable, and robust predictive models based on machine learning, contributing deeper insights to the current field.

## Acknowledgements

We thank all participants and investigators involved in the data producing including TCGA, CGGA, SRA database, and GEO database.

---

**Supplementary Figure S1. Evaluation of prognostic models constructed by Mime.**

**A.** C-index of StepCox[forward]-Ridge combined model across 9 cohorts. **B**. Expression level of genes involved in StepCox[forward]-Ridge combined model between control and PIEZO1 knockdown condition in G508 and G532. **C**. 3-year AUC of each model among different cohorts sorted by the average of AUC in validate cohorts. Black font of number mean that risk score calculated by this model predicted better outcome in corresponding cohort, otherwise predicted worse outcome. **D**. Same as C but for 5-year AUC. **E**. ROC curves of StepCox[forward]-Ridge combined model in different cohorts.

**Supplementary Figure S2. Comparison of AUC with previously established models.**

**A.** 3-year AUC predicted by StepCox[forward]-Ridge combined model and 95 published models across 9 cohorts. **B**. Same as A but for 5-year AUC.

**Supplementary Table S1. Basic information of nine included series.**

**Supplementary Table S2. Information about 18 cohorts accepting immunotherapy.**

**Supplementary Table S3. The characteristics of signature and their respective coefficients from previous studies used for comparison.**

**Supplementary Table S4. Information about 13 immunotherapy signatures used for comparison.**

## Key Points

1. We developed a flexible open-source R package, Mime, making it easier for users to construct machine learning-based integration models for a better analysis of specific signatures.
2. By using Mime, we constructed two excellent models based on PIEZO1-associated signature which presented high accuracy separately in predicting outcomes and immunotherapy response of patients compared with other published models.
3. Case on selection of SDC1 showed that Mime could efficiently identify critical genes for further exploration from a large number of features.

## Supplementary data

Supplementary data are available online.

## Ethics approval and consent to participate

Not applicable.

## Data availability

Data used to support our work are all public and data information can be acquired from the Supplementary Materials. Raw codes for package are available at GitHub (https://github.com/l-magnificence/Mime).

## Authors’ contributions

Study designing: HW-L, WZ, YH-Z, SY-WG and XJ-L. Data acquisition and analyzing: HW-L, WZ, YH-Z, AA-A, LH-D, ZY-P, WL, KP and DO-F. Manuscript writing: HW-L, WZ, YHZ and AA-A. Manuscript editing: SY-WG and XJ-L.

