## Supplementary figures and images for "Mime: A flexible machine-learning framework to construct and visualize models for clinical characteristics prediction and feature selection"

### Supplemental Figure S1

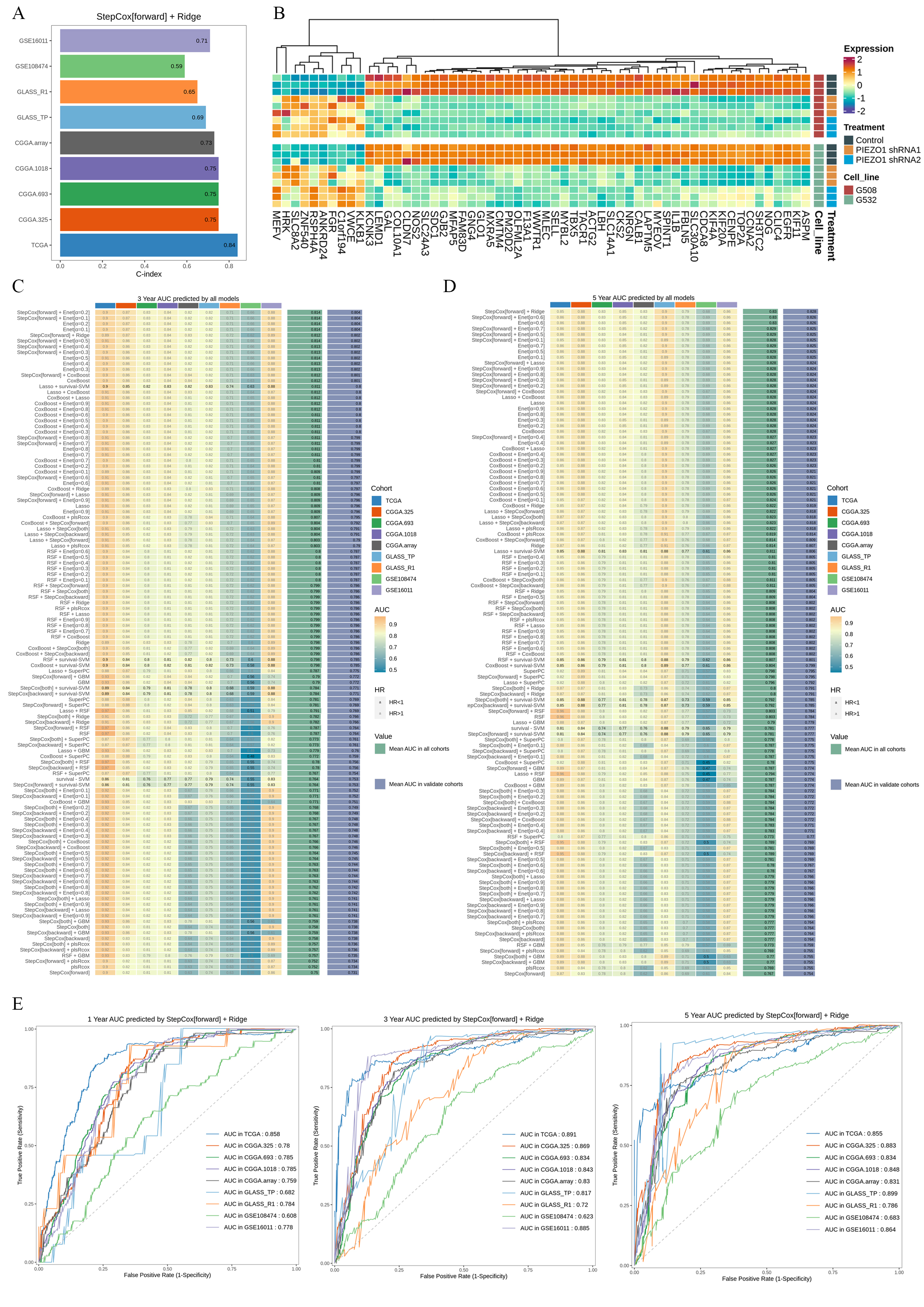

### Supplemental Figure S2

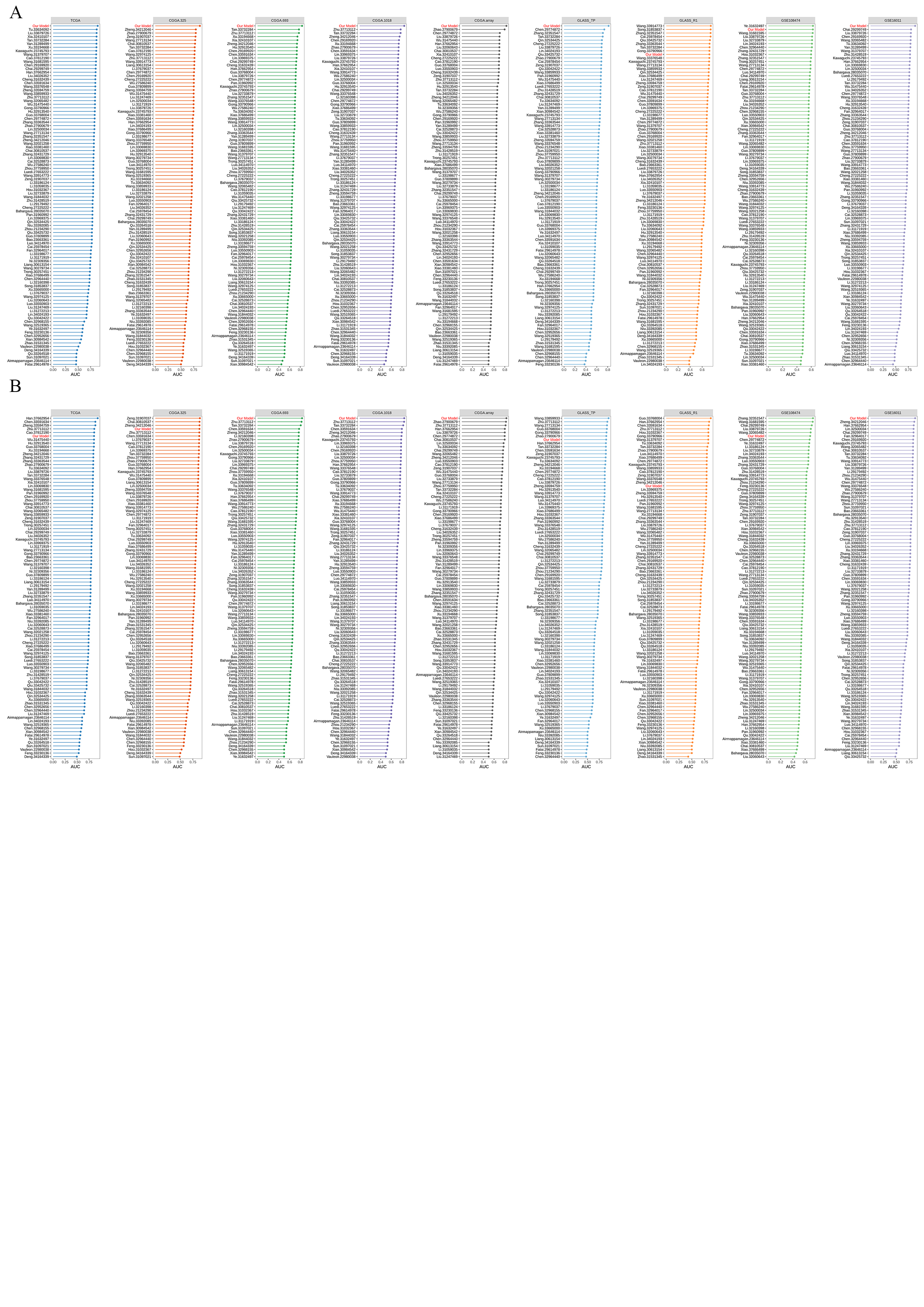
